# The main activatory and tension-sensitive transitions occur within Prozac sensitive down-states of the potassium selective TREK-2 channel

**DOI:** 10.1101/2020.10.22.351205

**Authors:** Michael V. Clausen, Jakob Ulstrup, Hanne Poulsen, Poul Nissen

## Abstract

The two-pore domain potassium selective (K2P) ion-channels TREK-1, TREK-2, and TRAAK essential mechanical stimulation sensors, and TREK-1/2 also targets for the antidepressant Nor-fluoxetine (Prozac). They respond directly to membrane tension by moving from the “down” to “up” conformation, a transition that is associated with a rise in open-probability. However, the mechanosensitive K2P (mK2P) channels can also open while occupying the down conformation, and although these channels are mostly closed, all structural models represent seemingly open conformations. To understand the dynamics between open/closed and up/down states and determine how membrane tension influences transitions between specific conformations, we use a novel method to analyze tension-driven activation of single purified and reconstituted TREK-2 channels. We screen a panel of prospective schemes to find the mechanism that best accounts for specific TREK-2 characteristics as tension-driven activation, suppression by Nor-fluoxetine, and single-channel kinetics.

To adequately describe TREK-2 behavior, mechanistic schemes require two separate tension-sensitive transitions, one that occurs between distinct down conformations and one that moves the channel between down and up states. As membrane tension activates TREK-2, it is a transition within the structural down conformations that account for the major increase in open-probability (> 100 fold); the move from down to up further promotes channel opening, but with much lower potency (~3 fold activation).

## Introduction

The mechanosensitive two-pore domain potassium selective (K2P) channels TREK-1, TREK-2, and TRAAK, set and regulate the negative membrane potential of, e.g., cardiac myocytes [1] and neurons [2], and underlie the repolarization at nodes of Ranvier during action potential propagation [3, 4]. As these channels are involved in mechanical events like cardiac arrhythmias [5], the sensation of pain [6, 7], and bladder control [8], and since axon membranes stretch during action potentials [9, 10], there is considerable interest in understanding how membrane tension activates mechanosensitive K2P (mK2P) channels.

As TREK-1 [11, 12], TREK-2 [13], and TRAAK [12] maintain their sensitivity to membrane stretch after purification and reconstitution into vesicles of defined composition, their mechanosensitivity derives directly from membrane tension. Crystal structures of TREK-2 [14] and TRAAK [15, 16] fall into two distinct classes: up- and down-conformations, of which the down-conformation occupies less volume in the membrane bilayer and has a larger intracellular part compared to the up-conformation (Fig. 1). TREK-1 structures have so far all been in the up-conformation [17].

**Figure 1.**
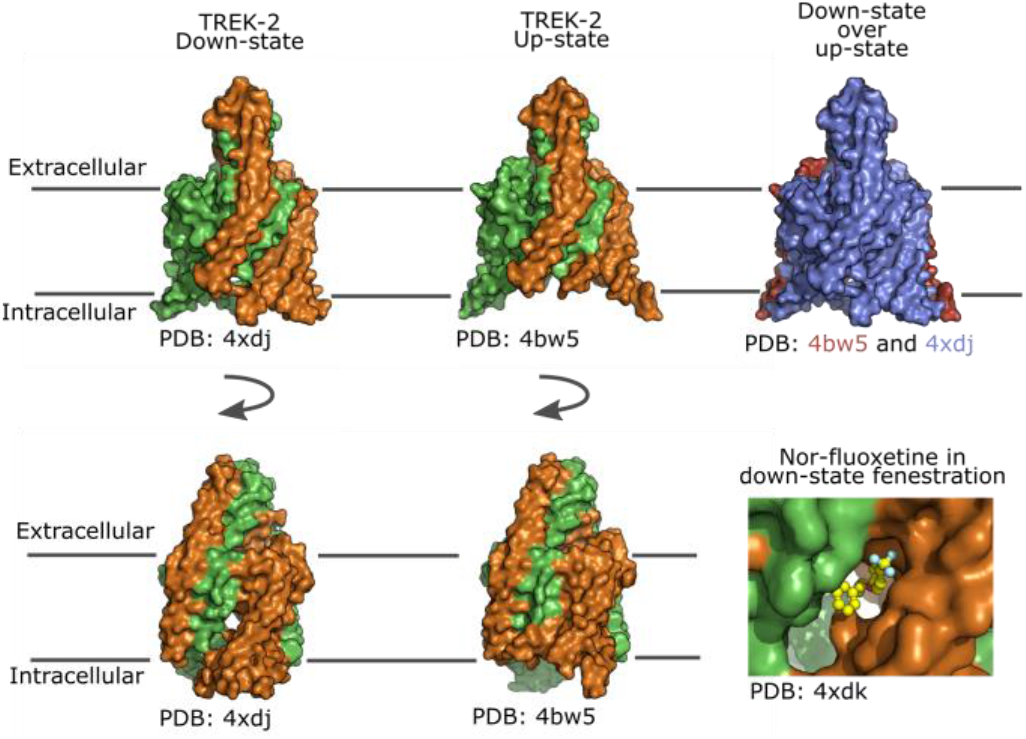
TREK-2 structural states and nor-fluoxetine binding. From the left, the figure shows two sides of TREK-2 in the down-state with individual subunits in green and orange; notice the transmembrane fenestration visible in the bottom panel perspective. In the middle, TREK-2 in the up-state is illustrated from the same views as the down-states; notice the absence of a transmembrane fenestration. In the top-right panel, the down-state structure (blue) is superimposed on the up-state (red) to show the relative expansion of the up-state within the membrane. The bottom right panel focus on the nor-fluoxetine (yellow) binding site within the down-state fenestration.

None of the 20 current structures of TREK-1, TREK-2, and TRAAK provides a clear insight into the channel gating mechanism because, in all of them, the conduction pathway appears similar and open. Substantial evidence suggests that the up-conformation has a higher open-probability (P_O_) [14, 15, 18] and that tension increases channel occupancy of the state [13, 18, 19]. From a theoretical perspective, we expect that a channel activated by membrane tension will expand within the membrane as tension increases [20], and cysteine bridges that lock TRAAK in up-conformations activate the channel [15]. A different line of evidence involves the antidepressant Fluoxetine (Prozac) and its metabolite Nor-fluoxetine (NF), which suppress currents through TREK-2 by binding within a transmembrane fenestration that is only formed when the channel is in a down-conformation [14] (Fig. 1). This state-dependent suppression, combined with the observation that stretch-activated TREK-2 channels are less sensitive to NF [14], argues strongly for a model where up-conformations have higher open-probabilities than down-conformations, and increased membrane tension favors the up-conformation. However, two highly active TRAAK mutants crystalize in down-conformations [16], and studies of TREK-2 show that some modes of activation, e.g. extracellular acidification and Rb^+^-ions, operate via down-conformations [18].

Most potassium selective channels, e.g., the voltage-gated (Kv) and inwardly rectifying (Kir) channels, have a lower intracellular gate at the helix bundle crossing and an upper gate that involves the selectivity filter; mK2P channels in contrast control conductance exclusively at the selectivity filter [21, 22], and since all TREK-1, TREK-2, and TRAAK crystal structures have the same filter conformations, the mechanistic coupling between up/down transitions and channel opening/closing events remains elusive.

Here, we exploit a global single-channel fitting procedure [23] to screen different mechanistic schemes that describe TREK-2 activation by membrane tension and state-dependent inhibition by NF. With an unsupervised approach, we compare 19 schemes and find that a single mechanism successfully accounts for TREK-2 activity. Our final model shows that membrane tension activates TREK-2 through two consecutive transitions; the first is between different down-conformations, and this accounts for a larger part of channel activation, while the second, smaller contribution, is the transition from a down- to an up-confirmation.

## Methods

### Protein production and reconstitution

Human TREK-2 (KCNK10) with N- and C-terminal truncations (Gly^67^ to Glu^340^) [14] was cloned into the pEG vector [24] and used for transfection of HEK293-F cells. Cells were harvested 72 hours after transfection and resuspended in a lysis buffer consisting of 50 mM HEPES pH 7.5, 150 mM NaCl, and 1 mM PMSF and then lysed using a Sonopuls sonicator. Membranes were isolated from the lysate by ultracentrifugation and then solubilized in 50 mM HEPES pH 7.5, 200 mM KCl, 1% OGNG, and 0.1% CHS. The solubilized sample was mixed with Strep-Tactin^®^ resin (IBA Lifesciences) and washed with 50 mM HEPES pH 7.5, 200 mM KCl, 0.18 % OGNG, and 0.018 % CHS. The strep-tag was cleaved off overnight at 4°C using the 3C-protease. The protein sample was purified further on a Superose 6 with a running buffer consisting of 20 mM HEPES pH 7.5, 200 mM KCl, 0.12% OGNG, and 0.012% CHS. Peak fractions were saved at −80°C and thawed before use.

Giant Unilamellar Vesicles (GUVs) were made from 1,2-diphytanoyl-sn-glycero-3-phosphocholine by electroformation in 1 M sorbitol using Vesicle Prep Pro (Nanion Technologies). GUVs were destabilized with detergent before purified TREK-2 was added to a final concentration of ~1-4 ug/ml. Detergents were removed by incubation overnight at 4⁰ C with Bio-Beads (Bio-Rad).

### Single-channel recording and idealization

All electrophysiological recordings used the Port-a-Patch planar patch-clamp system (Nanion Technologies) connected to an Axopatch 200B amplifier via a Digidata 1550B1 (Molecular Devices). Giga-Ohm seals formed as TREK-2 containing GUVs ruptured on contact with the chip glass surface over a microstructured aperture [25]. All data was recorded with a 100 kHz filter and a sample rate of 500 kHz. Additional digital filtering at 20 kHz, kept the number of false-positive openings detected during idealization below one per minute (Fig. S1A). We used a BK4052 waveform generator (B&K Precision) to deliver square-pulses of specific lengths through the patch-clamp rig and used a plot of observed vs. applied pulse lengths to define a 20 μs deadtime used throughout the study (Fig. S1B).

We used a set of relations that take total recording time and mean P_O_ to estimate the probability that a patch with only single openings have two or more channels [26]; the probability that patches in this study had two or more channels is less than 10^−10^. Single-channel data was idealized in MATLAB using a half-amplitude threshold-crossing algorithm, and a 20 μs deadtime was imposed (Fig. S2).

### Mechanistic schemes

The goal is to derive a discrete state Markov model that accounts for TREK-2 activity. Throughout this paper, we use the term scheme when referring to a non-parameterized model, i.e., a scheme containing information only about how specific states are connected and no information about the rates at which the channel transits between distinct states. The mechanistic schemes used through this study rely on three assumptions:

Assumption 1) Channels can occupy several distinct structural states; transition rates between these states are either tension-sensitive or tension-insensitive. The justification is that while some channels have P_O_’s that are affected by membrane tension, others still move between distinct states, but in a manner that is unaffected by tension [12].
Assumption 2: TREK-2 can exist in several distinct structural sub-states that belong to one of two principal conformations, an up-state (NF insensitive) and a down-state (NF sensitive) [14, 18].
Assumption 3: Volume expansions within the membrane underlie tension-driven channel regulation [20], and we assume that transitions that involve volume changes are tension-sensitive, even if they originate from conformations that are tension-insensitive per se (Fig. S3). It follows that transitions between up- and down-states are tension-sensitive as the up-state occupies additional intramembrane volume [14, 15, 19].

### Fitting of single-channel data to mechanistic schemes

To probe prospective schemes, we use a four-stage *fit-protocol* that integrates information from dwell-time data obtained at three different tension levels into a model with fixed and tension-sensitive rates (Fig. S4). The *fit-protocol* uses three fitting scripts based on the SCAIM procedure [23] that uses the methods derived by D. Colquhoun, A.G. Hawkes, and A. Jalali to correct for missed events [27, 28] (available at https://github.com/biologichael/SCAIM). The three fitting scripts comprise i) a *single-fit* that finds optimal parameters to an input scheme and a dwell-time segment, ii) a *multi-fit* that fits three dwell-time segments to a scheme while keeping specific parameters fixed and others free, and iii) a *simplex-optimization* routine that optimizes initially seeded parameters.

### Model-based simulation and evaluation

Each parameterized scheme obtained after stage 2 and 4 of the *fit-protocol* was used for stochastic simulation of single-channel data imposed with a 20 μs deadtime. All schemes have states that are either up (NF insensitive) or down (NF sensitive). To evaluate how single-channel behavior of specific models change upon application of NF, two additional closed states, representing one and two NF molecules bound, were added to each down-state. The NF off-rate for these states was set to 1000 s^−1^ and data simulated based on mid-P_O_ models with NF on-rates set to 0, 1000, and 2000 s^−1^. Data were simulated in triplets for each tension-setting and NF level and evaluated against both specific values from the observed data, e.g., P_O_ and burst-length, and general trends observed throughout observed data, e.g., burst-lengths that decrease with NF and inter-burst closure lengths that gets briefer with tension. Positive evaluations required that each of the three simulated sets for each condition fulfilled the given criteria.

### Term definitions

Scheme: a scheme that specifies how specific states are connected and include information about the functional status of each state, i.e., open/closed and up/down. Schemes also separate transition rates into tension-insensitive and divide tension-sensitive into expansions and contractions.

Model: a model is a parameterized scheme with both fixed and tension-sensitive rates. The term model-mode refers to a model with specified tension-sensitive rates.

Segment: a segment is a single stretch of alternating channel openings and closures that have a constant open-probability throughout.

Fit-protocol: the term *fit-protocol* refers to the procedure illustrated in Fig. 4S.

Multi-fit: refer to a specific fitting procedure used in stage 3 of the *fit-protocol*.

Set: a set refers to a collection of three dwell-time segments representing low, mid, and high open-probabilities.

## Results

### Channel open probability as a proxy for membrane tension

Purified human TREK-2 was reconstituted in 1,2-diphytanoyl-sn-glycero-3-phosphocholine. For patches with a single TREK-2 channel, membrane stretch was delivered by suction protocols ranging from −40 to −120 mbar, demonstrating that channel P_O_ increases when the pressure decreases (Fig. 2 A and B). The effect is partly reversible since the release of the suction decreases P_O_. Still, the membrane stretch can have a longer effect, altering channel baseline activity, i.e., P_O_ in the absence of suction (Fig. 2 A before and after - 120 mbar suction jump). Between and during suction protocols, channel activity remains stable (Fig. 2 C), but equal suction does not always produce a similar change in P_O_ (Fig. 2 B).

**Figure 2.**
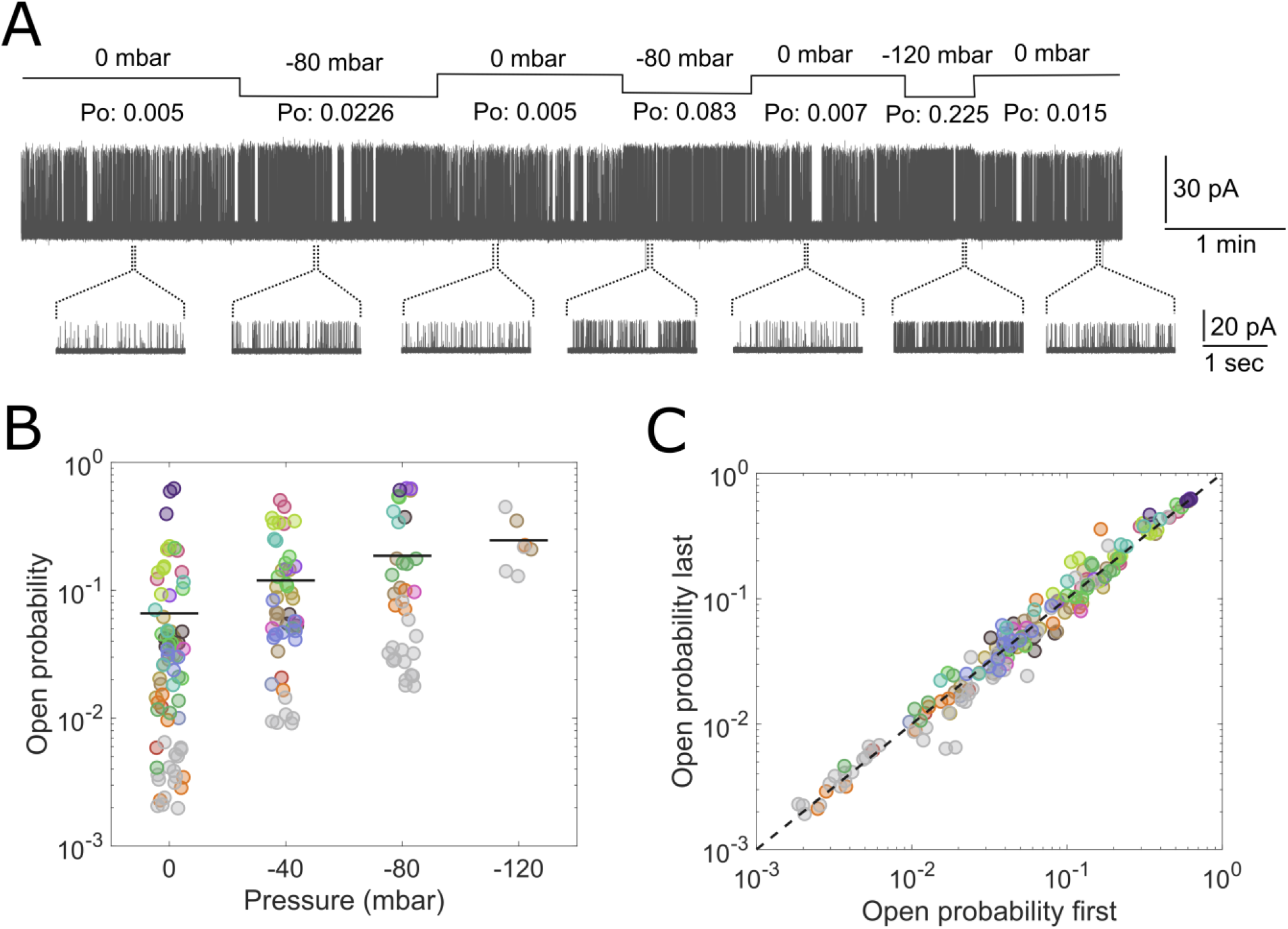
Channel activity changes following stimulation via membrane stretch. A) Nine minutes of single-channel data obtained with suction settings and associated P_O_’s indicated above the trace, and expanded views of single-channel openings (upward deflections) below. B) P_O_’s from 170 segments of single-channel data from 16 patches plotted as a jittering scatterplot against the holding pressure. Data from individual patches are color-coded, and the mean P_O_ for each pressure setting is given as a horizontal line. C) For each segment, dwell-times separated into equally sized vectors representing the first and last half of the entire segment, are plotted against each other.

In this study, we use data from 16 single-channel patches from which 184 segments (170 at distinct pressure levels, and 14 from NF trials) with stable P_O_’s were isolated and used for analysis.

Because of the simple experimental system with a single channel in a membrane of a single lipid species, we assume that the only parameter that influences channel activity is membrane-tension. We cannot quantify membrane tension; only modulate it with suction, but we see that the Po correlates with the pressure (Fig. 2B). Therefore, to examine the mechanism underlying tension-driven channel activation, we study single-channel kinetics at different P_O_’s.

### Single-channel parameters underlying tension-driven activity changes

The single-channel kinetics can be determined from analyses of the dwell-times observed in open and closed conformations. The relative abundance of the open and closed conformations determines the P_0_, and we, therefore, analyze them at different P_0_’s. For the closed states, the analysis of dwell-time histograms shows that at least six components, c1-c6, are required to describe the distributions (Fig. 3A). For the higher P_0_’s, it is clear that a subset of closed states are destabilized (Fig. 3A): Both dwell-time length and total occupancy diminish for the three longer closed components (c4, 5 and 6), the briefer (c1, 2 and 3) are less affected and remain <2 ms in all segments.

**Figure 3.**
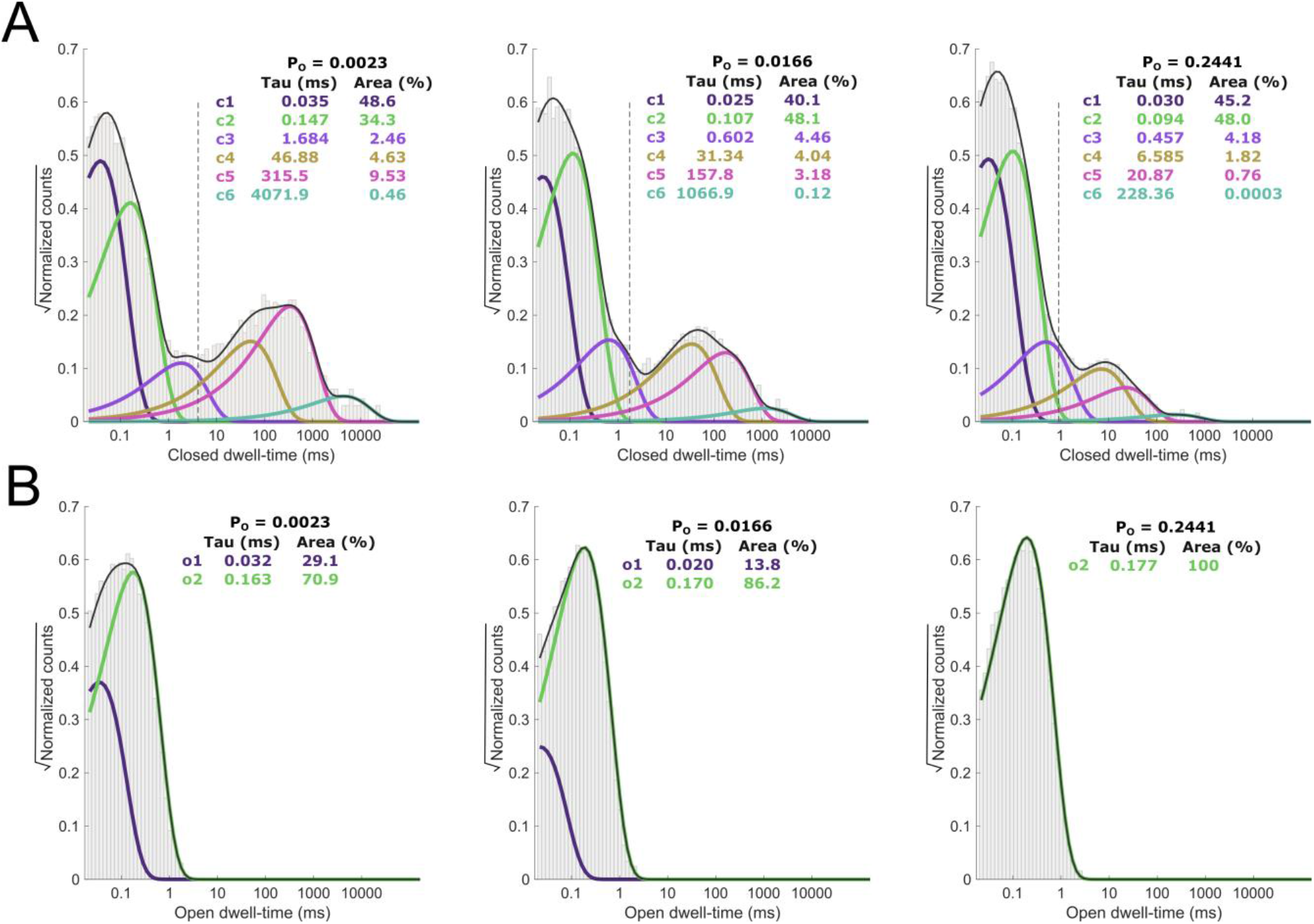
Maximum likelihood fits to open and closed dwell-time distributions. Maximum likelihood fits to open and closed dwell-time distributions obtained at different segments from the same patch. A) Closed dwell-time distributions of low P_o_ segments require six components (c1 through c6) to produce proper fits. Fewer components may adequately describe higher P_o_ segments, but we maintain six to simplify comparison as P_o_ increase. The vertical dotted line between components c3 and c4 indicates the position of the critical closed time t_crit_ used to define bursts of openings. B) Fits to open dwell-time distributions require two components (o1 and o2) when P_o_ is low; in high P_o_ segments, however, the o1 component completely disappears.

Dwell-time histograms of open-times also change with tension (Fig. 3B). In the low P_O_ segment, fits to open dwell-time histograms comprise two components (o1 and o2), and as tension increases, the brief component o1 disappears, and the extended o2 tend to increase in length (Fig. 3B).

To further characterize our single-channel data, we define a critical closed time (t_crit_) [29] for each segment by numerically solving

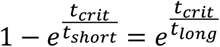

where t_short_ is the longer of the brief components (c3), and t_long_ is the briefest of the long components (c4) (Fig. 3A). We characterize groups of more than two openings flanked by closures longer than t_crit_ as bursts of openings (Fig. 4A) and find a strong correlation between channel P_O_ and burst length (Fig. 4C). Similarly, we find that the mean duration of closures between bursts (closures longer than t_crit_) decrease with P_O_ (Fig. 4D).

**Figure 4.**
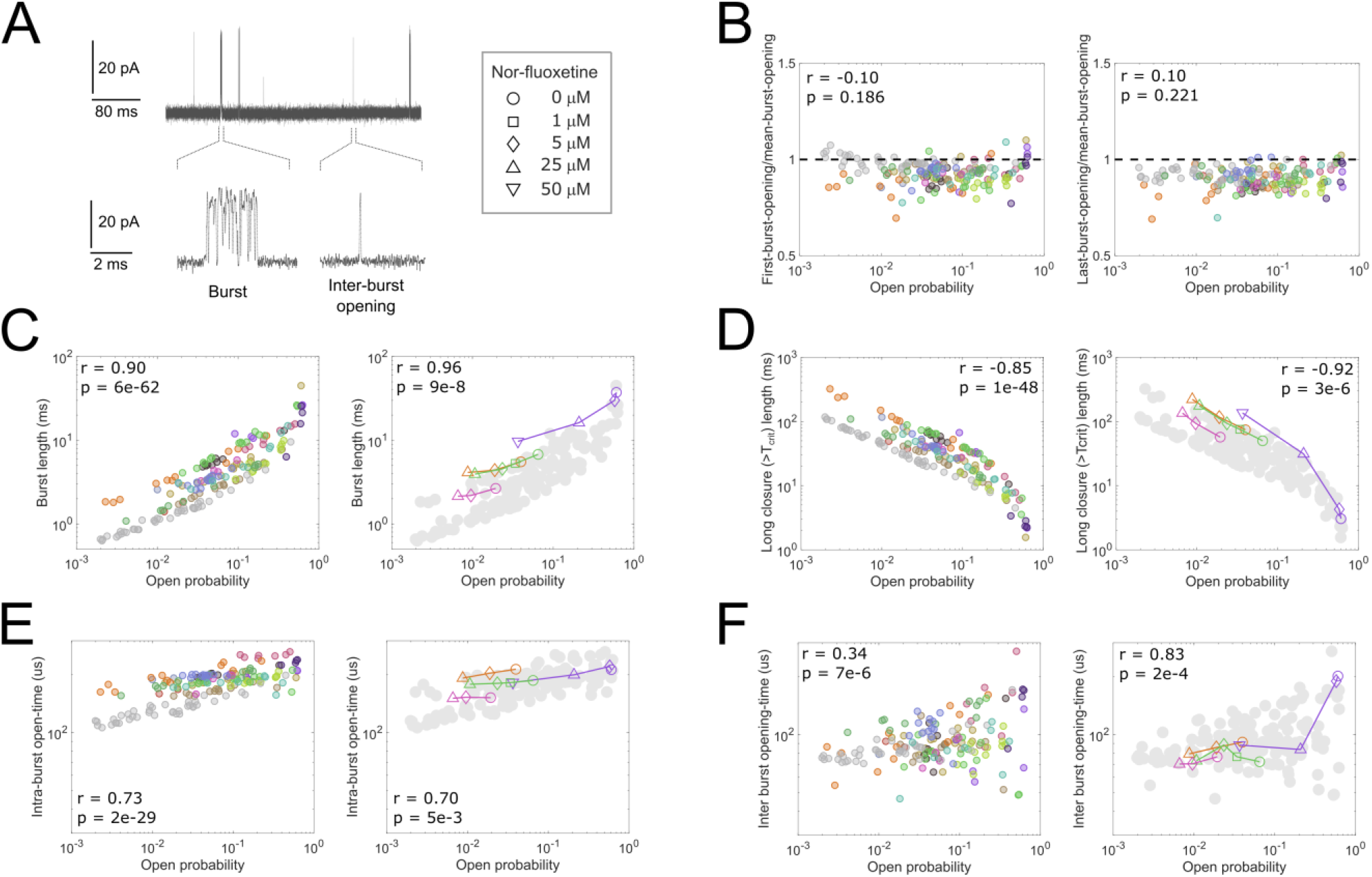
Single-channel characteristics. A) Excerpt from single-channel recording illustrating a typical burst with openings separated by closures < t_crit_ and a single opening flanked by closures > t_crit_. The inserted table with nor-fluoxetine (NF) concentrations refer to panels C, D, E, and F. B) Individual bursts of every segment analyzed by dividing the lengths of the first (left panel) and last (right panel) opening by the mean open-time of that burst. Each marker gives the mean value of the ratio for individual segments plotted against the P_O_; the dotted line indicates a ratio of one. C) Mean burst lengths of discrete segments plotted against P_O_ with the left-hand panel representing points from data that differ only by the level of membrane tension. In the right-hand panel, four experiments with NF concentrations, as indicated by the table in (A), overlay gray-toned data from the left panel. Individual NF experiments are connected with lines and color-coded. The strength of correlation given by the Pearson coefficient r along with a p-value for null hypothesis tests is given in both panels for tension and NF data, respectively. D) As (C) but with data on the Y-axis representing mean lengths of closures > t_crit_. E) As (C) but with data on Y-axis representing mean open times within bursts. F) As (C) but with Y-axis demonstrating mean lengths of open-times outside bursts.

For the open channel states, plotting the length of the first and last opening of a burst divided by the mean open-length of that burst versus P_O_ shows that bursts begin and end with brief openings (average 93% and 90%, respectively), and this is similar for the different P_0_s, suggesting that the phenomenon is tension-independent (Fig. 4B). Because mean open-times within bursts increase with tension (Fig. 4E), the lengths of first and last burst openings thus increase similarly. In Fig. 4F, we examine the relationship between lengths of single openings outside bursts and P_O_ and find that these are briefer than intra-burst openings (Fig. 4E). As P_O_ increases, the mean length of inter-burst openings shows higher variation, an observation that occurs concomitantly with a decrease in the frequency of single-openings (single openings per second fall with increasing P_O_, r = −0.31, pval = 3e-5).

In four patches, we examined the effect of NF on single-channel characteristics and found that the antidepressant decreases burst lengths (Fig. 4C), increases the mean closed time between bursts (Fig. 4D), and reduces the mean open-time within bursts (Fig. 4E).

### A novel strategy to evaluate putative mechanistic schemes

Our principal goal is to find a model that encompasses what we know about TREK-2. Therefore, we need a mechanistic scheme that specifies how distinct open and closed states are connected, and that categorizes specific states into the two major conformational classes (up and down). Furthermore, a scheme must have tension-sensitive rates to describe transitions that involve volume expansion/contraction within the membrane (assumption 3).

With a basis in a global fitting procedure that finds model parameters for a scheme without any prior seeding, we designed a four-stage fit protocol that incorporates information from three different dwell-time segments (Figure S4). The purpose of the complicated procedure is to limit the parameter-space toward models that can account for a wide range of open-probabilities by adjusting only pre-defined tension-sensitive rates. The underlying assumption behind our strategy is that the success of a proposed scheme in reproducing various features of the observed data correlates with the similarity between the actual underlying mechanism and the input scheme. We test this assumption in figure S5 by simulating three sets of data using a defined model with four variable rates. To approach the real scenario, we impose a 20 μs deadtime on the simulated data before proceeding with the fitting-protocol.

Figure S5C tests how eight different schemes produce fits that can simulate data that resemble the observed. The three schemes that most successfully fulfill the evaluation criteria also correspond best to the true model. Scheme_Z has a structure identical to the underlying model, Scheme_Z2 has a similar construction with four closed and two open states that, via two variable transitions, connect with an open and two closed states. Finally, Scheme_Z7 reveals a limitation in the approach; while the scheme is a truncated version of the true scheme, it provides a better evaluation. Compared with Scheme_Z, Scheme_Z7 lacks a closed state (C9) that has kinetics similar to state C8 (on-rates 12000 s^−1^ vs. 5000 s^−1^ and off-rates 5 s^−1^ vs. 15 s^−1^). A more focused search-space with fewer parameters gives Scheme_Z7 a better evaluation. Still, stricter criteria, e.g., comparisons of closed dwell-time histograms, could ultimately show that Scheme_Z represents the optimal mechanism.

### Models require two separate tension-sensitive transitions

Figure 5 demonstrates the evaluation of sixteen schemes following 150 independent fits divided between three sets that each includes three dwell-time segments (Table S1). In schemes A through H, tension-sensitive rates are placed exclusively between down- and up-conformations; schemes I through P have additional tension-sensitive rates dividing either up-states (schemes J, L, and N) or down-states (schemes I, K, M, O, and P) into subpopulations of states.

**Figure 5.**
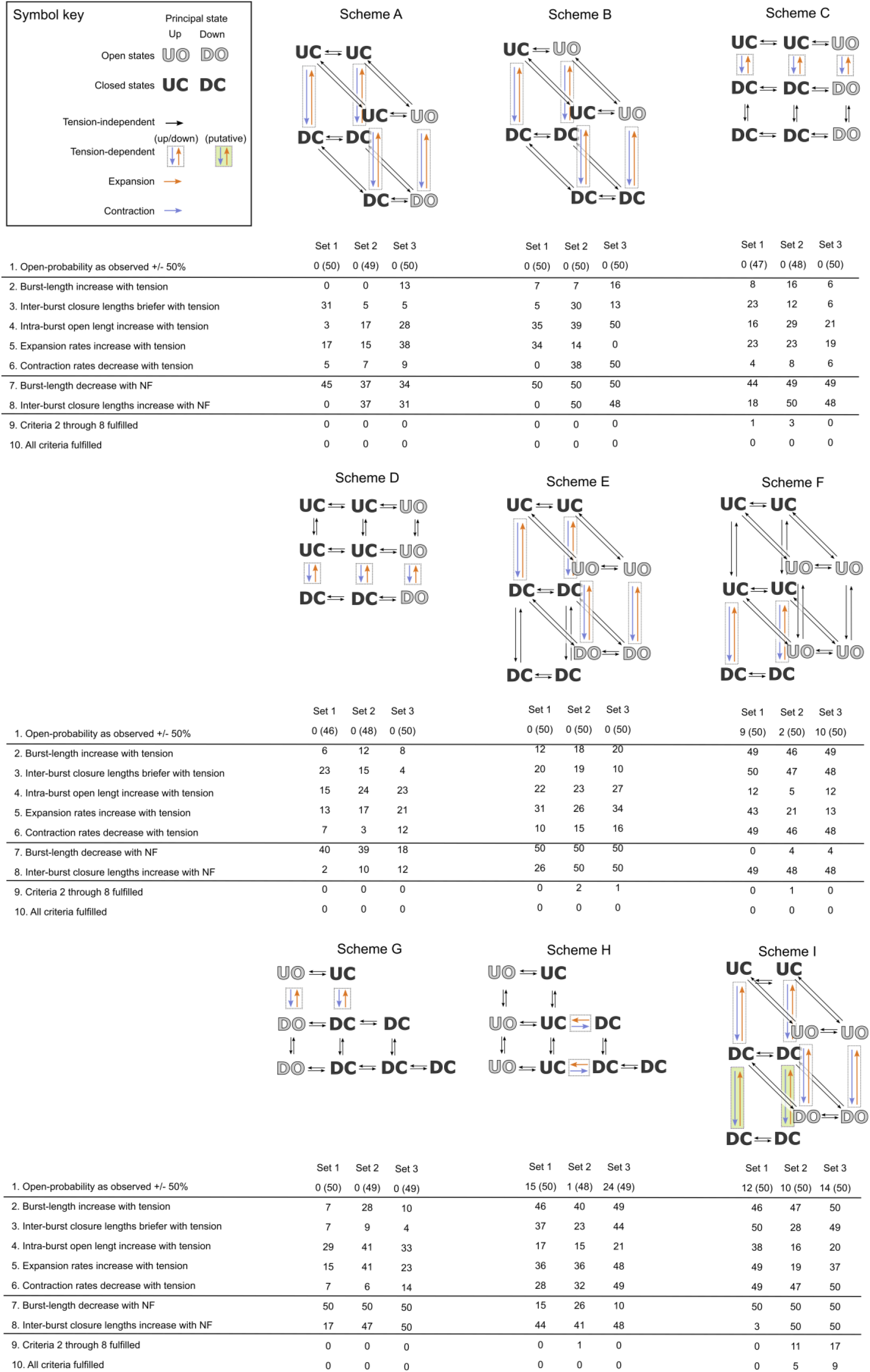
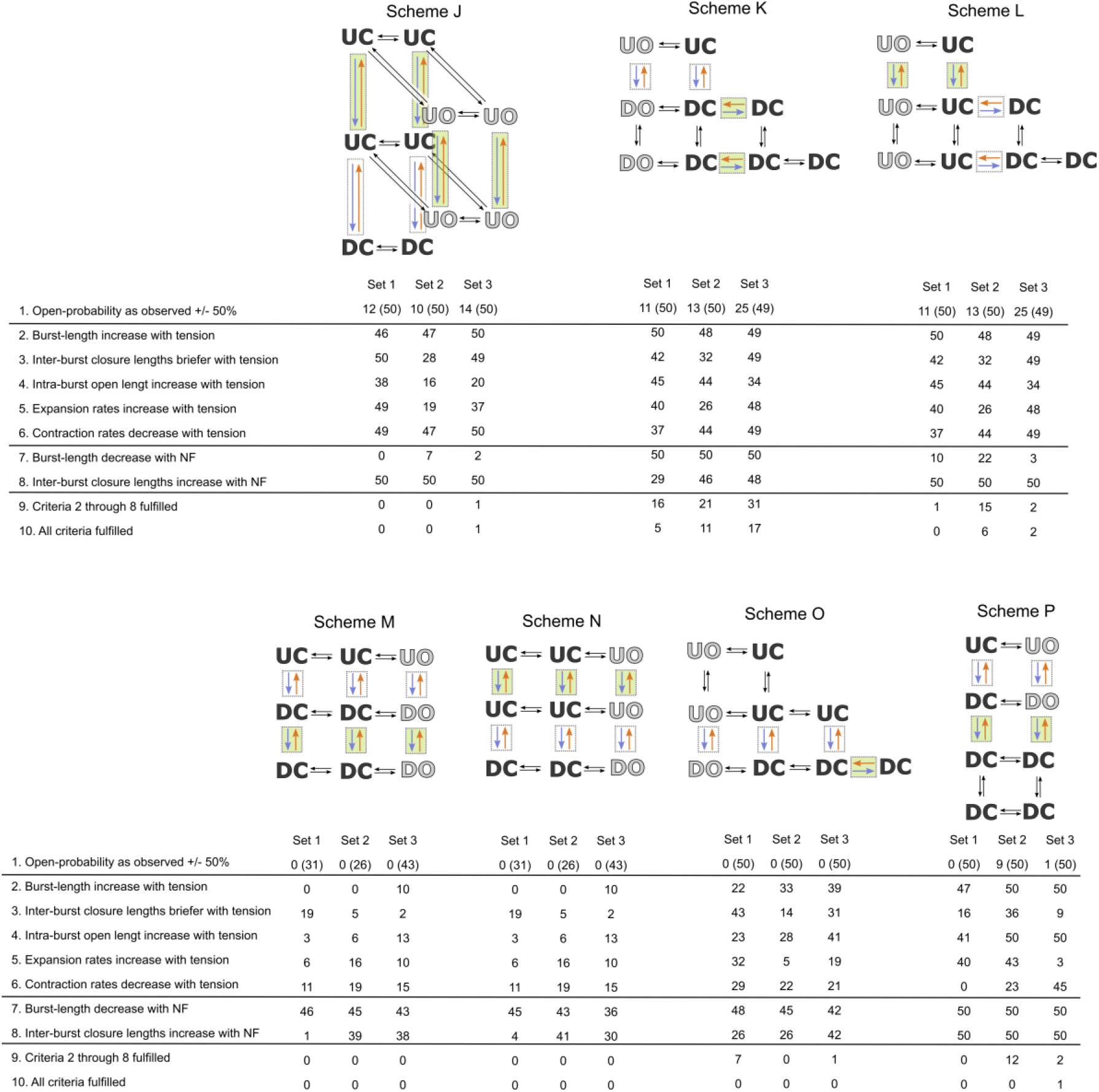
Initial evaluation of mechanistic schemes. Sixteen schemes specify connections between open (gray/O) and closed (black/C) states that are in up (U) or down (D) conformations. Transitions are either tension-insensitive (black arrows) or tension-sensitive (rates within dotted boxes), and tension-sensitive rates are expansions (orange arrows) or contractions (blue arrows). Schemes I to P includes putative tension-sensitive transitions beyond the move between up and down highlighted as green boxes. Each scheme is assessed after fitting to three sets (Set 1, Set 2, and Set 3) of dwell-times, that each includes three dwell-time segments representing low, mid, and high open-probabilities (table S1 and fig. S4). Fifty fits per set through stages 1 and 2 of the *fit-protocol* (Fig. S4), are evaluated below each scheme through simulation of data per the derived parameters. Evaluation criteria 1 tests if simulated data reproduce the P_O_ of the three dwell-time segments of the given set, with the number in parenthesis giving successful fits to the mid-P_O_ segment alone. Evaluation criteria 2 through 6 examine if burst-length, inter-burst closures, intra-burst open-times, expansion- and contraction-rates increase or decrease going from low to mid to high P_O_ fits. Evaluation criteria 7 and 8 examine the simulated effect on burst lengths and inter-burst closure lengths as increasing concentrations of NF bind to down states. Evaluation criteria 9 count the number of fits that successfully meet criteria 2 through 8, while criteria 10 counts those that fulfill all criteria.

We divide tension-sensitive rates into contraction-rates and expansion-rates that, in accordance with criteria 5 and 6, must either decrease and increase with tension. To focus the search-space toward solutions that fulfill these criteria stages 1 through 3 of the *fit-protocol* use constraints that fix rates characterized as expansions or contractions to stay equal. This maneuver limits the number of parameters that must change in a defined direction between low, medium, and high P_O_ segments to two, namely one for expansions and one for contractions, but also puts a considerable limitation on our models. The constriction biases the fitting procedure toward models where all tension-sensitive transitions are structurally related, and while stage 4 releases the constraint, this bias remains an integral part of the models. However, without fixed expansion and contraction rates, no fits comply with criteria 5 and 6.

In Figure 5, the numbers in parenthesis under evaluation criteria 1 validate that most stage 1 fits successfully reproduce the P_O_ of the mid-segment. However, following consecutive fits to low- and high-P_O_ segment data (stage 2), where only tension-sensitive parameters remain free, only a subset of fits effectively matches the open-probabilities of the input dwell-time segments.

Most schemes can meet criteria 2 through 8 separately, and schemes A, B, D, G, M, and N all demonstrate examples where these criteria are met collectively (criteria 9). Only schemes with states divided into three subpopulations separated by tension-sensitive rates demonstrate examples where all criteria are meet (schemes I, J, K, L, and P). Of these, the additional tension-sensitive transition works better when located within the down-states (scheme I vs. J and K vs. L). Among the sixteen schemes, Scheme_K stands out as markedly better within the three sets of dwell-times and remains the only scheme that successfully produces models for set 1.

### Single-openings outside bursts originate from specific down-conformation

Of the sixteen schemes, only Scheme_K successfully fits all sets (Fig. 5). Each successful fit (criteria 10) provides three sets of parameters for Scheme_K and hence three model-modes that account for TREK-2 activity: one mode for each of the three dwell-time segments of the given set. Figure S6 shows plots of the factors that expansion- and contraction-rates change with between low-, mid- and high-P_O_ segment model modes.

The multi-fit protocol in stage 3 (Fig. S4) uses a global search strategy to optimize a set of parameters by simultaneously fitting the three dwell-time segments of a set. The same set of parameters, with tension-sensitive rates varying by factors determined after stage 2 (see above), are evaluated on the three dwell-time segments, and the fit-value of the algorithm [23] identified as the sum of these values. Therefore, the multi-fit protocol does not favor any of the dwell-time segments. The procedure in stages 1 and 2, on the other hand, produce results that match better with the mid-P_O_ dataset.

Figure 6 demonstrates evaluations of Scheme_K and three offshoots with sub-indices a, b, and c, following stage 4 of the *fit-protocol* (Fig. S4). The nine initial evaluation criteria are as in figure 5, and criteria 1 illustrates that the multi-fit protocol is less poised toward mid-P_O_ segments than the single-fit approach. The evaluation in figure 6 includes five additional measures that specifically address characteristics of single-channel behavior (criteria 10-14). Scheme_K fails to produce models with all criteria fulfilled for the set 1 dwell-times and performs poorly on the set 3 segments. For both sets, especially criteria 10 and 11 cause problems, and as these involve channel openings, we constructed and evaluated three derivatives of Scheme_K, each with an additional open state.

**Figure 6.**
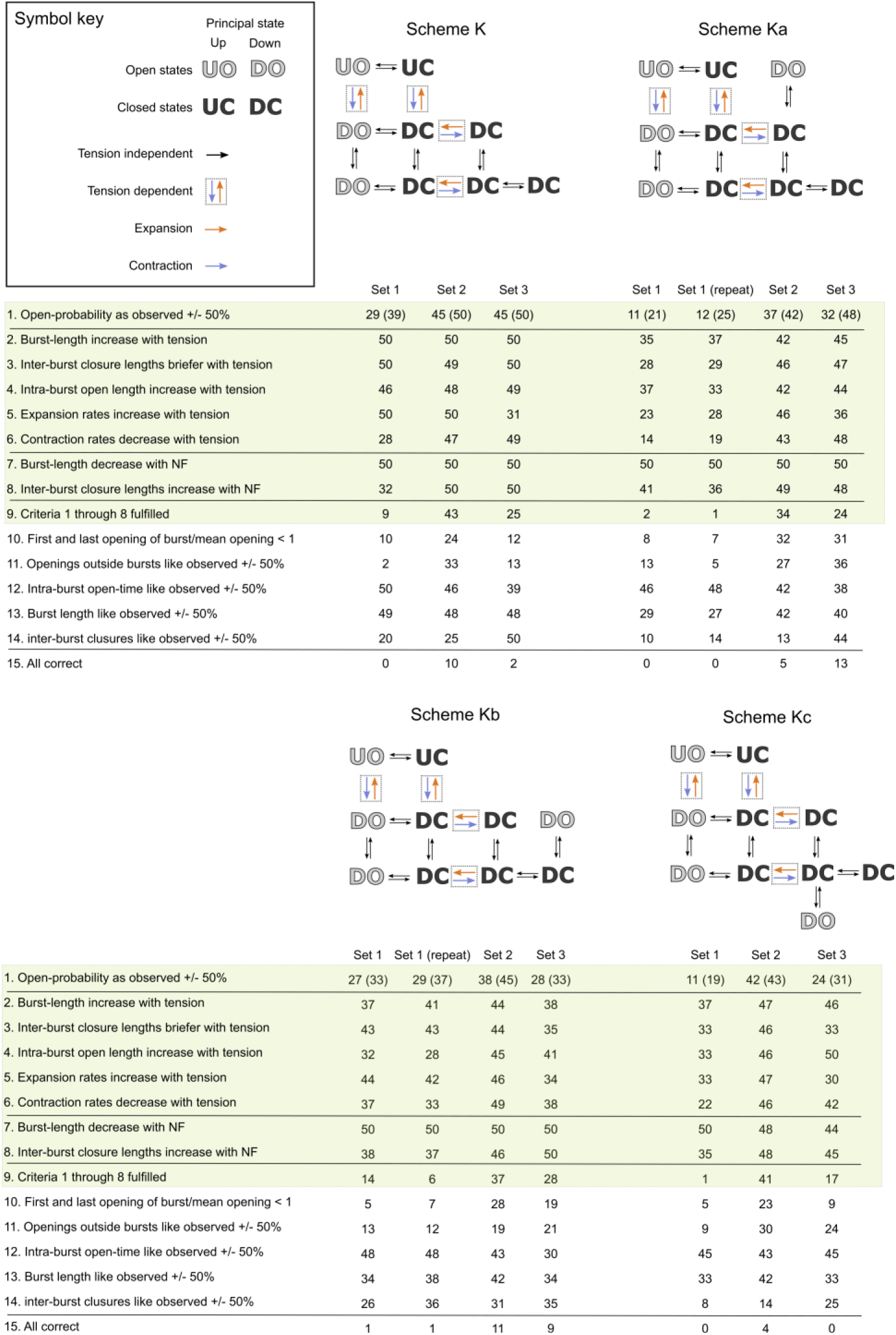
Optimization toward a final model. Four schemes evaluated after fitting through the four stages of the *fit-protocol* (Fig. S4) using data from three sets, each including segments representing low-, medium- and high-P_O_ (table S1). One hundred fifty repeats of the *fit-protocol* for each scheme, fifty for each set, give 150 models with parameters to match the three input segments. Therefore, each model comes with three modes, and the evaluation compares data simulated from each model-mode to observed data from the three input segments; only simulated data that match all tension-levels counts toward a positive evaluation. The exceptions are criteria 1 and 11: under criteria 1 the evaluation of mid-P_O_ data is shown alone in parenthesis, and under criteria 11 no comparisons are made between high-P_O_ data as single openings are nearly absent. The entire procedure was repeated for set 1 for schemes Ka and Kb, as indicated by the added column.

The impact of an additional open state depends on how it connects with Scheme_K; when linked as in Scheme_Kc, the resulting evaluation is inferior to Scheme_K, whereas linkage as in Scheme_Kb shows substantial improvements (figure 6). As compared to Scheme_Ka, Scheme_Kb mainly stands out for its ability to handle the data of set 1. While Scheme_Kb better evaluates most criteria for fits to set 1 data than Scheme_Ka, only a single set model successfully accounts for all criteria. To verify these results, we repeated the entire fitting procedure for the set 1 segments and find that Scheme_Kb provides a better mechanism. Besides, the repetition of the set demonstrates the constancy of the process.

### A model that explains mechanical activation of TREK-2

To identify common features in the 22 Scheme_Kb fits that meet all criteria, we plot all parameters from each mid-P_O_ fit in the same graph (Fig. S7A). Some parameters are strikingly similar in all fits (e.g., i2, i5, and i11); others have values spread out (e.g., i4, i6, and i19). It is worth noting that for dispersing parameters, the equilibrium constants between the states they connect show similar trends: i6/i4 is >3 (22 of 22), i19/i21 is between 0.5 and 2 (20 of 22), i24/i22 is >3 (18 of 22), and i7/i15 is >3 (20 of 22).

As a consensus model, we selected the fit with parameters closest to the median values (Fig. S7B).

In our TREK-2 model, tension-sensitive rates divide channel conformations into three groups, one with up-states, one with mid-down states, and one with down-states (Fig. 7A). Based on the selected model parameters, we calculate the P_O_ of the three groups and find that the major increase in P_O_ occurs within down-states, where P_O_ increases almost a thousand-fold from deep-down to mid-down. In contrast, it increases by less than four-fold from mid-down to the up-states. This tendency is shared among the 22 Scheme_Kb fits that fulfill all criteria: In these models, P_O_ increases from down to mid-down by more than 100-fold and from mid-down to up by less than 6-fold.

**Figure 7:**
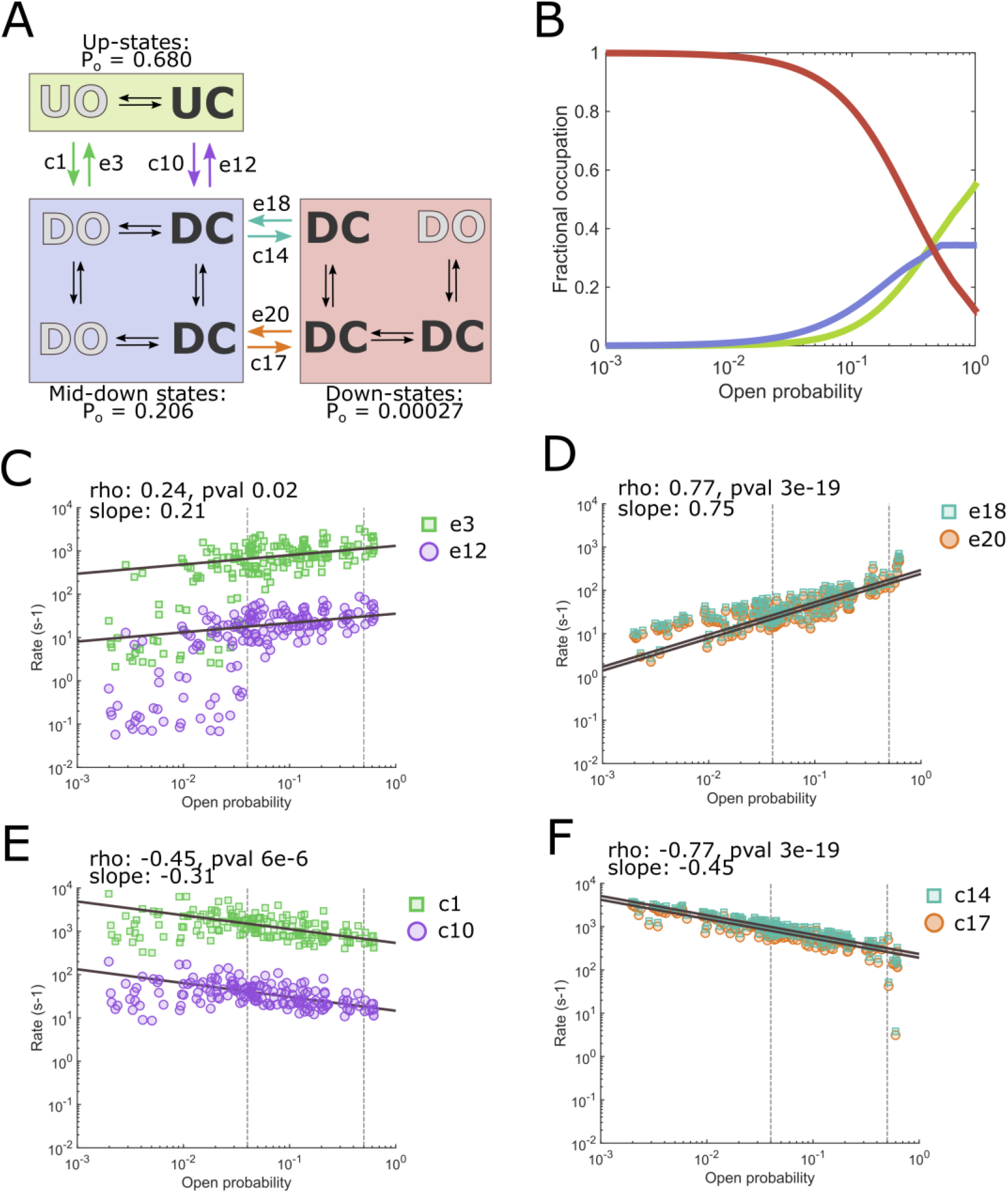
Sequential activation. A) Mechanistic scheme with transition rates for tension-sensitive expansions and contractions. Tension-sensitive rates separate the scheme into three principal states, down, mid-down and up, as indicated by colored boxes. The P_O_ for each principal state is given along each box. B) The plot show how the relative time spent in each of the principal states (colored as in panel A) change as tension increase P_O_. Data for the plot is calculated with tension-sensitive rates as extrapolated in panels C-F and tension-insensitive parameters as deducted in Fig S7B. C-F) Parameters for the eight tension-sensitive transition-rates obtained from model fitting to the 170 dwell-time segments, with tension-insensitive rates fixed and tension-sensitive seeded as in fig. S7B. For each panel the strength of the linear correlation is given as Pearson correlation coefficient (rho) and the tension sensitivity as the linear regression slope, both values are derived from the area between the vertical dotted lines (P_O_ 0.04 to 0.5), and the regression line extrapolated to cover the entire data span.

We are not quantifying membrane tension in our analysis; therefore, we cannot infer information about tension-sensitivity in terms of which tension-levels have the most impact on activity. Nevertheless, the steepness of the relationship between Po and a tension-sensitive rate still directly reflects how instrumental the rate is in changing channel activity as tension increases.

To examine tension-sensitivity of down to mid-down and mid-down to up transitions, we fitted our model to the 170 dwell-time segments with fixed tension-insensitive rates and tension-sensitive seeded as in the table of fig S7B. The relative difference between expansions and contractions were fixed to maintain microscopic reversibility [30]. Figure 7C-F shows plots of expansion and contraction rates derived from these fits. The spread in mid-down to up transitions at low Po (Fig 7C and E) and down to mid-down at high Po (Fig 7 F) reflects a dominance of down-states at low Po and an absence at high Po. Therefore, to compare tension sensitivity, we focus on the stretch between P_O_ 0.04 and 0.5, where there is apparent linearity without outliers.

The Pearson Correlation Coefficient shows that the expansion rates from mid-down to up-states (Fig. 7C) increase with P_O_, but that the correlation is weak; in the other direction, the correlation is more robust and the tension-sensitivity steeper (Fig. 7E). The transition between mid-down and up is >30 fold faster when the channel is open than when it is closed (e3 and c1 compared to e12 and c10), a tendency that is shared among the majority of the Scheme_Kb fits that fulfill all criteria; here 17 of 22 have e3 and c1 rates faster than e12 and c10.

Between down and mid-down states, the positive and negative correlations are strong (Fig. 7D and F), and the tension-sensitivity is more pronounced than for mid-down/up transitions. While the particular closed state that the channel occupies during transition is without influence (e18 vs. e20 and c14 vs. c17), the tension-sensitivity between the forward and reverse reaction is markedly stronger for the expansion from down to mid-down (slope 0.75 vs. slope −0.45).

With rates for tension-sensitive transitions derived from linear regression fits as in figure 7C-F, we calculate the fractional occupancy of the three principal states (Fig. 7B). The plot demonstrates that down-states are dominating at low P_O_, and up-states become the preferred conformations only at P_O_ >0.4; this supports the interpretation that the spread in the data at low P_O_’s in figures 7C and E is due to an almost complete absence of transitions to up-states.

## DISCUSSION

Ion-channels move dynamically between states that are conductive or closed. Distinct conformations can differ in their interaction, both with chemical ligands like drugs, lipids, and neurotransmitters, and with the physical environment like temperature and membrane-tension. Models that describe and predict ion-channel behavior can help us understand how and why cells, tissues, and organisms react to a given stimulus and may suggest new rational drug design strategies. However, to make such models, we must assign particular states that a channel can occupy and how the ion-channel moves between these states under specific conditions. The model should be as simple as possible and yet complex enough to explain observations without making assumptions that are not physically meaningful.

Here, we exploit a novel strategy to identify a model that accounts for the single-channel behavior of TREK-2 while respecting prior knowledge regarding how the channel moves from NF-sensitive down states to NF-insensitive up-states as membrane tension increases.

Because the only input we feed into the algorithm is idealized data and prospective schemes, the entire evaluation is unbiased toward specific mechanisms. In this light, it is striking that among sixteen schemes, all with six closed states and between two and four open conformations, only one manages to fulfill all evaluation criteria for the three sets of input dwell-times (figure 5). The flexibility to accommodate all selection criteria relates more to how distinct states are connected than to the number of free parameters; Scheme_K has 13, Scheme_P has 11, and Scheme_C has 16. This is also evident in figure 6, where the position of an additional open state influences whether the evaluation improves or worsens.

The analysis in figure 5 also demonstrates the advantage of a scheme with a sequential activation mechanism. To account for TREK-2 activity with a P_O_ that ranges from <0.005 to >0.5, schemes with tension-sensitive rates simply between down- and up-states must have down- and up-states with P_O_’s of <0.005 and >0.5. This requirement conflicts with the reduced burst lengths triggered by NF binding because burst activity terminates as channels enter down-states, which would limit the ability for NF to affect burst lengths. This obstacle is complicated further by the tension-sensitivity of intra-burst open times; if down-states account for only a negligible part of burst activity, then the effect on open-time by prolonged time in up-states becomes undetectable. Of eight schemes with tension-sensitive rates only between down- and up-states, only two produce fits that span the P_O_ of the input segments (Scheme_F and Scheme_H); these at the same time rank bottom in predicting a reduced burst length upon NF binding.

Schemes with sequential activation mechanisms, where tension-sensitive transitions divide them into three sections, deal much better with the evaluation criteria; of eight schemes with sequential activation, five demonstrate examples of fits that fulfill all criteria.

A sequential activation mechanism fits well with *in silico* observations. Here, a stretch of a membrane with TREK-2 in the down-conformation triggers multiple structural rearrangements that occur in sequence. The final transition is an upward movement of transmembrane helix 4 (TM4), which closes the fenestration that constitutes the binding site for NF and brings the channel into the up-conformation [19].

The notion that down- and up-conformations represent principal structural states with different levels of activity comes from three concurrent papers. TRAAK mutants with cysteines situated to lock the channel in the up-conformation are activated by oxidizing conditions and lose mechanosensitivity [15]. In a different study, however, two TRAAK mutants with elevated basal activity and amputated sensitivity to mechanical stimulation crystalized in the down-conformation [16]. Finally, TREK-2 crystalizes in both up- and down-conformations, and only down-states comprise a binding site for the activity inhibitor NF [14]. The finding that the potency of NF decreases in TREK-2 activated by mechanical stimulation supports the idea that tension increases channel open-probability by stabilizing up-conformations [14].

Our model for TREK-2 activity and mechanical activation unifies the conclusions above because it explains both why locking the channel in the up-state increases activity and diminishes mechanosensitivity, why NF potency decreases with membrane tension, and finally, how mutations that stabilize specific down-states can increase basal activity and reduce tension-sensitivity. Therefore, our study supports the conclusions based on state-dependent blockage with NF, namely that TREK-2 can gate in the down-, as well as in the up-conformation [18]. A similar conclusion was reached in a molecular dynamics simulation experiment where two meta-stable down conformations with high open probabilities were observed [31].

Our model does not only support prior findings; it contributes new insight into how TREK-2 operates. We find that the major tension-sensitive and activatory transitions occur within channel down-states, namely between down- and mid-down-states. This predicts that a channel hindered in reaching the up-conformations, e.g., due to a mutation, a physiological interaction, or pharmacological interference, will maintain substantial mechanosensitive properties if kinetics within down states are preserved.

It is remarkable that we, with several atomic resolution structures of TREK-1, TREK-2, and TRAAK, as well as a growing number of supporting *in silico* experiments, still lack a fundamental understanding of what occurs at the gate during channel opening and closure. One study suggests that TREK-2 closes as carbonyl S3 of the selectivity filter flips and that this occurs more often in down states [32]. Another study, also using computer simulations based on up-/down-conformations of TREK-2, finds a different filter gating mechanism, namely that the outer part of the selectivity filter obtains a non-conductive pinched conformation [31]. While our study does not provide direct information about which residues move to close the channel, our model does offer clues and predicts distinct functional states that will be useful in the dissection of the structural mechanism. We find that the energy barrier for the final upward movement of TM4 is smaller when the channel is open; this implies some steric interference between TM4-down/up and filter-open/closed transitions. Importantly, we find that most of the activation associated with increased membrane tension occurs before TM4 moves up and closes the fenestration.

Our finding that closure of the fenestration by TM4 plays only a minor part during activation by membrane tension raises questions concerning the physiological role of the maneuver. Besides the well-known interaction with NF and Fluoxetine, the transmembrane fenestration also serves as a docking site for lipid acyl chains [15, 19]. The mK2Ps are homodimers or heterodimers with individual subunits from TREK-1, TREK-2, or TRAAK, and therefore include two fenestrations [33–35]. Interestingly, a TREK-2 structure with brominated fluoxetine’s bound at the two fenestrations (PDB: 4xdl) has subunits with TM2, TM3, and TM4 situated differently (Fig. 8). This demonstrates that at least two structurally distinct TREK-2 down-states can bind fluoxetine. If different lipid acyl-chains interact in specific manners with subsets of mK2P down-states, then the fenestration may represent part of a gearing mechanism that sets the obtainable level of mechanical activation. In a membrane with lipids that interact weakly with the fenestration, mK2Ps can enter the up-conformation and activate completely. If, on the other hand, a membrane contains lipids that interact with the fenestration in both mid-down- and down-states, tension can still activate the channels, but to a lower extent. Finally, if a membrane has lipids that interact strongly with the down-state and prevent entry into mid-down states, mK2P’s will have a blunted response to mechanical stimulation.

**Figure 8:**
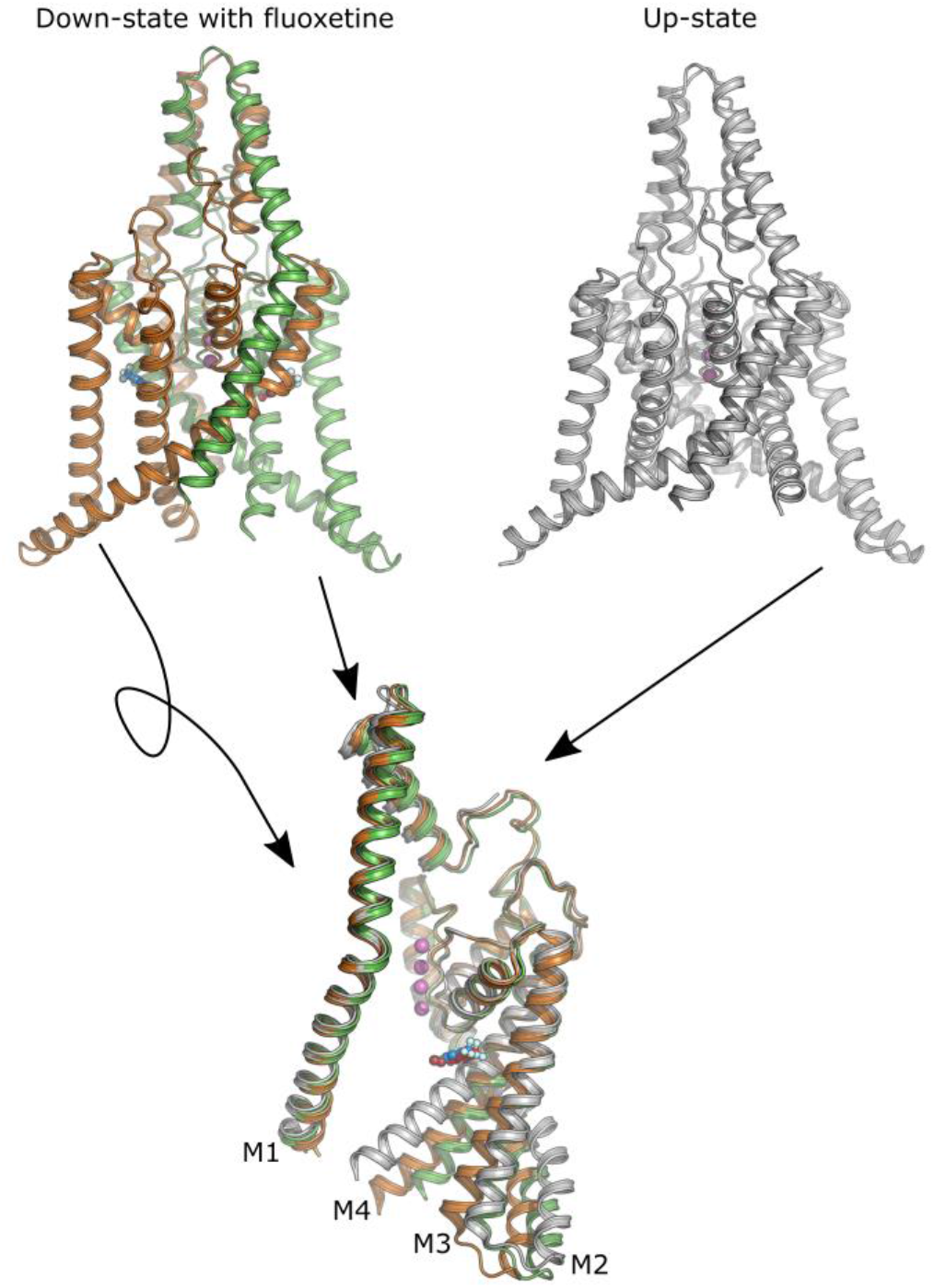
Three distinct structural positions of TREK-2. TREK-2 crystalized in the down-conformation with a Fluoxetine molecule bound (stick/sphere) in each of the two fenestrations (PDB: 4xdl), individual subunits colored green and orange, respectively, and three potassium ions in the filter shown as purple spheres. TREK-2 in the up-conformation shown with both subunits in light gray (PDB: 4bw5). The image bellow feature subunit 1 (orange) from the down-state flipped and overlayed with subunit 2 (green) and finally, a single corresponding subunit 2 of the up-state (white). The four transmembrane helices of the subunits are indicated as M1 through M4, and the subunits are superimposed with the selectivity filter as reference.

On-target toxicity refers to a classical problem in pharmacology, where a drug acting with high potency on its target results in adverse effects [36]. Our conclusion, that much of the mechanical activation of TREK-2, and possibly the related TREK-1 and TRAAK channels, occurs while the channels are in down-states, advocates for a strategy whereby we can dampen mechanical activation without completely abolishing it.

Finally, we develop and validate a novel approach to a difficult problem; with a global fitting strategy that corrects for missed events, we can screen distinct mechanistic schemes and identify those that best account for what we know about ion-channel activity and regulation. Models based on this method provide an excellent framework for additional functional and structural studies.

## ACKNOWLEDGMENTS

We are grateful to Jacob Lauwring Andersen for strategy development for channel overexpression and purification. M.V.C was supported by the AIAS-COFUND fellowship program under the EU’s FP7 for Research, Technological development and Demonstration (grant agreement no 609033) and the Aarhus University Research Foundation. Further project support was provided by a Lundbeck Foundation professorship grant to P.N. (R310-2018-3713) and an infrastructure grant from the Carlsberg Foundation (CF17-0910) to M.V.C. and P.N.

## AUTHOR CONTRIBUTIONS

M.V.C designed research, performed single-channel experiments, analyzed data, and wrote the manuscript. J.U. overexpressed and purified TREK-2 channels and commented on the manuscript. H.P. commented on the data analysis and manuscript. P.N. contributed project resources and commented on the data analysis and manuscript.

---

**Fig. S1:**
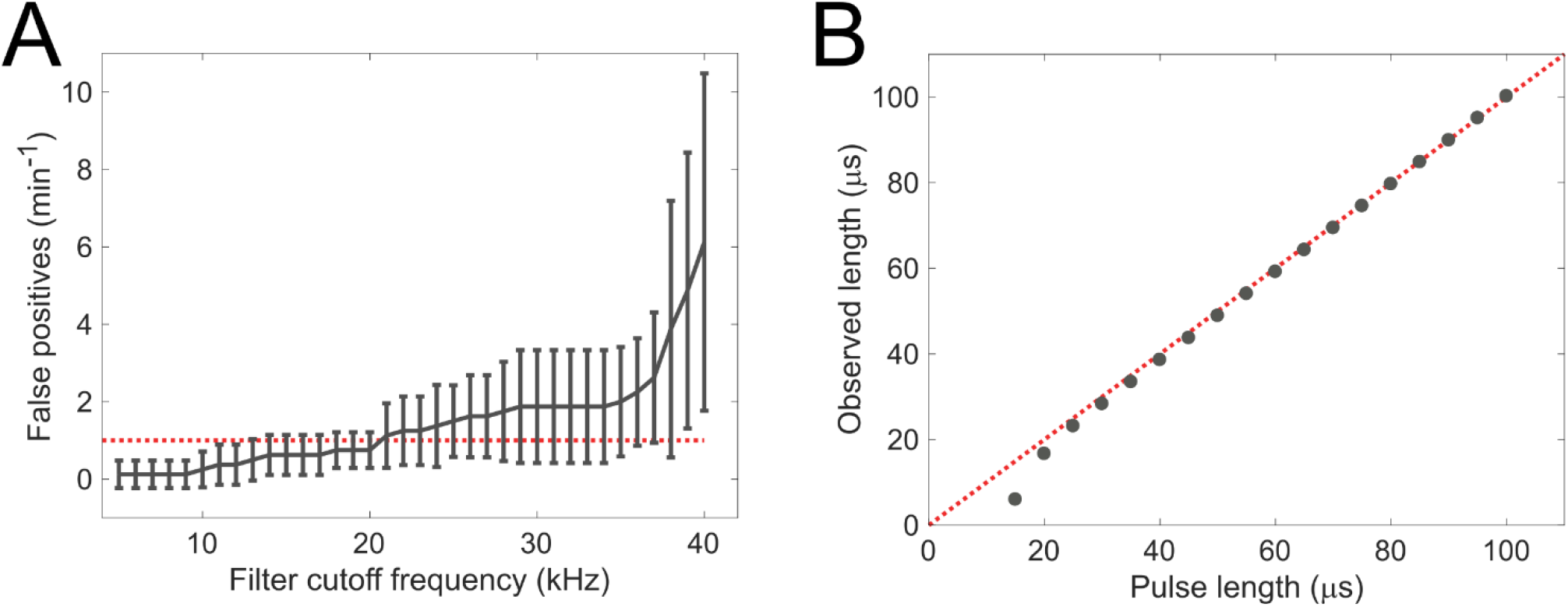
A) Recordings (> 1 min) with 100 kHz filtering and a 500 kHz sample rate from five empty patches held at 80 mV were further digitally filtered as given on the X-axis. False positives evaluated with a half-amplitude-threshold crossing algorithm with the threshold set at 12.5 pA. B) Square pulses with lengths as given on the X-axis were delivered to the patch-clamp rig with a function generator. After sampling and filtering as in A, observed event lengths were determined with half-amplitude-threshold crossing and given on the Y-axis.

**Fig. S2:**
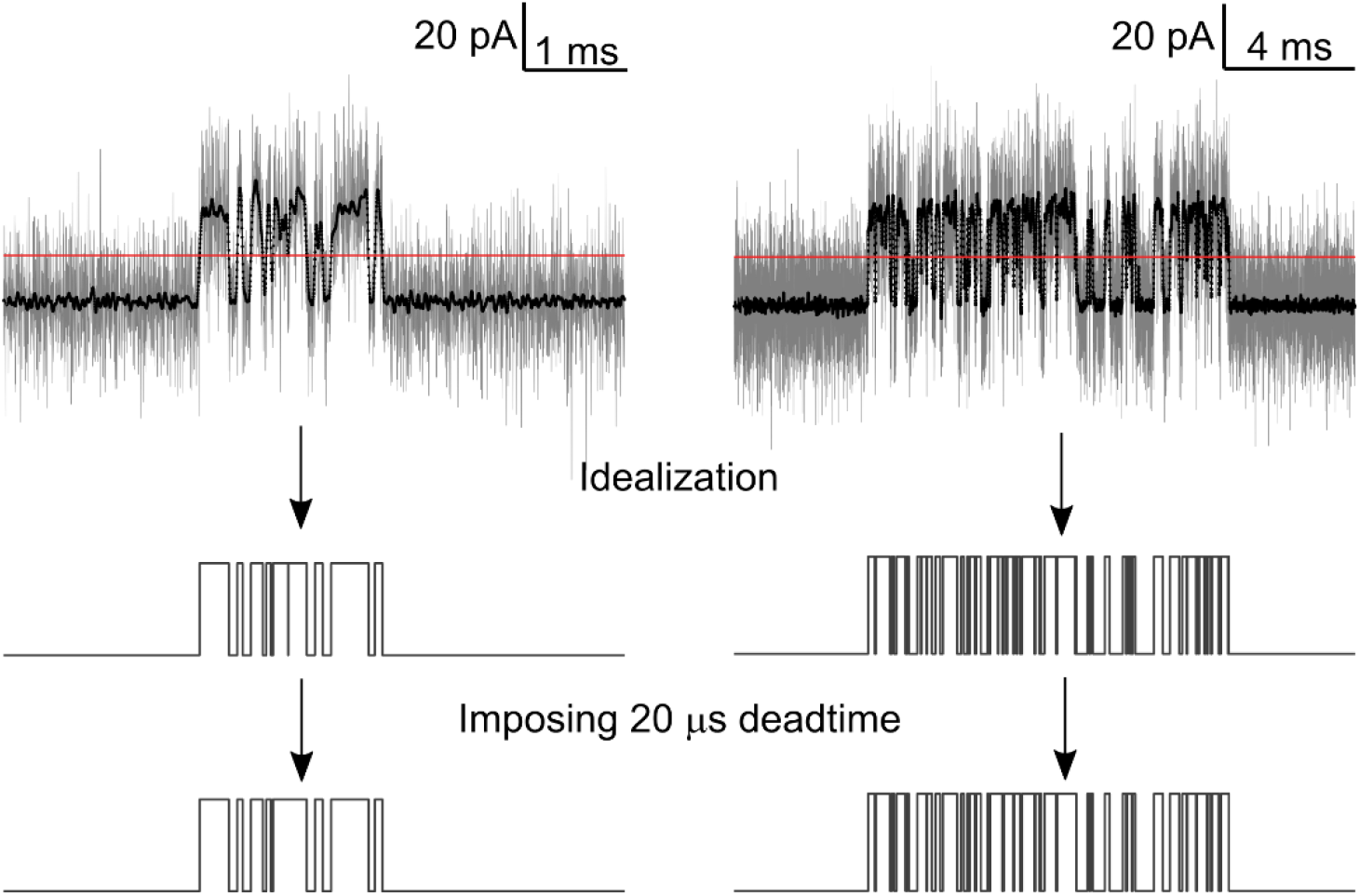
Two panels illustrating how raw data was processed to idealized data with well-defined time resolution. Single channel data sampled at 500 kHz with a 100 kHz filter in gray, the same trace following additional digital filtering at 20 kHz overlaid in black. The red line set the half-maximal amplitude threshold used during idealization. In the middle panel, show idealized data, the bottom panel idealized data after a 20 μs deadtime is imposed.

**Fig S3.**
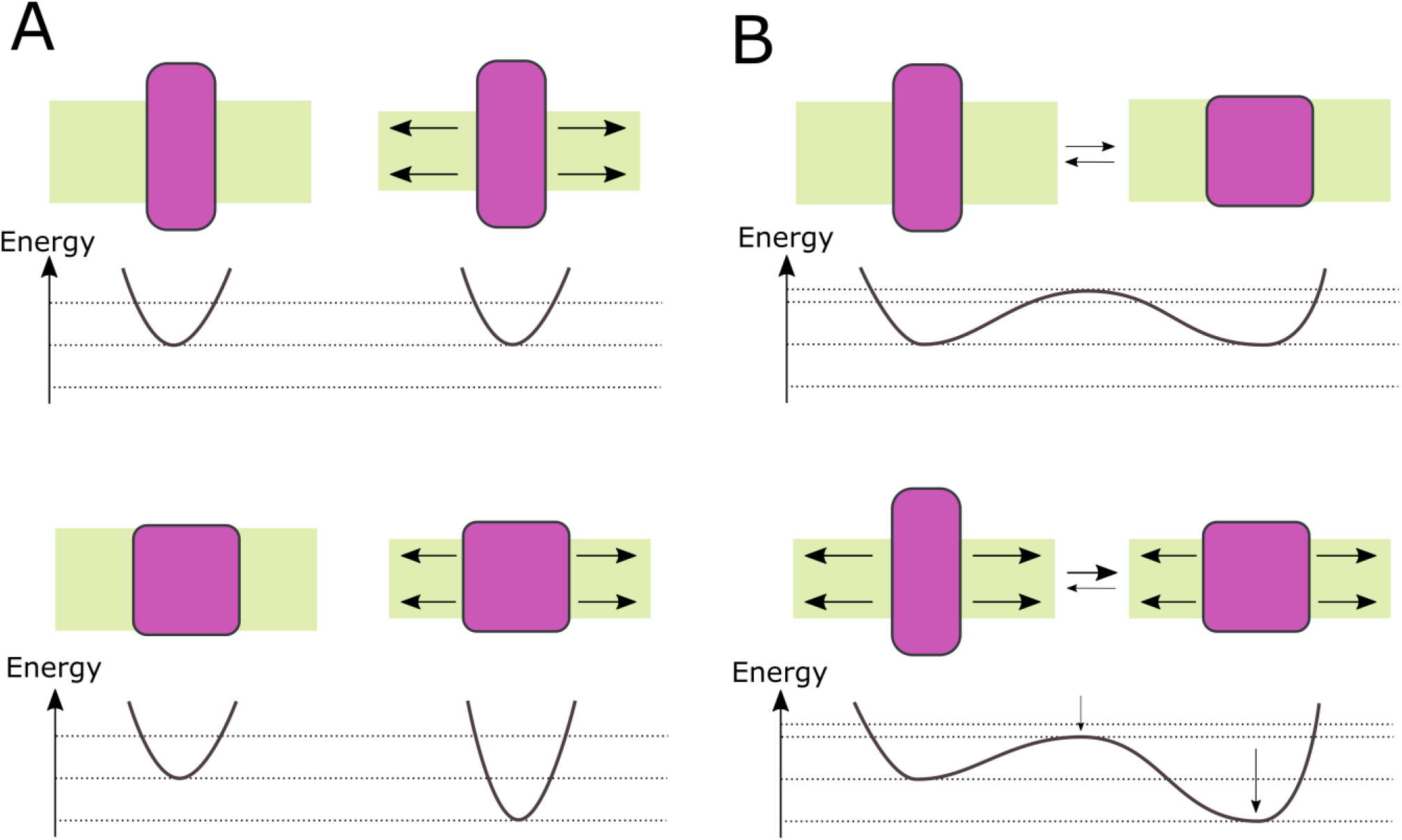
A) Two distinct channel conformations, one that is unaffected by membrane-tension (top panel), and one that fit better in a tense membrane (bottom panel). Increased membrane tension is illustrated with horizontal arrows in the membrane (green box) B) If the two states are connected, transitions between them are affected by membrane tension, both because of the structural stability of the tension-sensitive state change, and because the transition toward the tension-sensitive state involves volume expansion.

**Figure S4).**
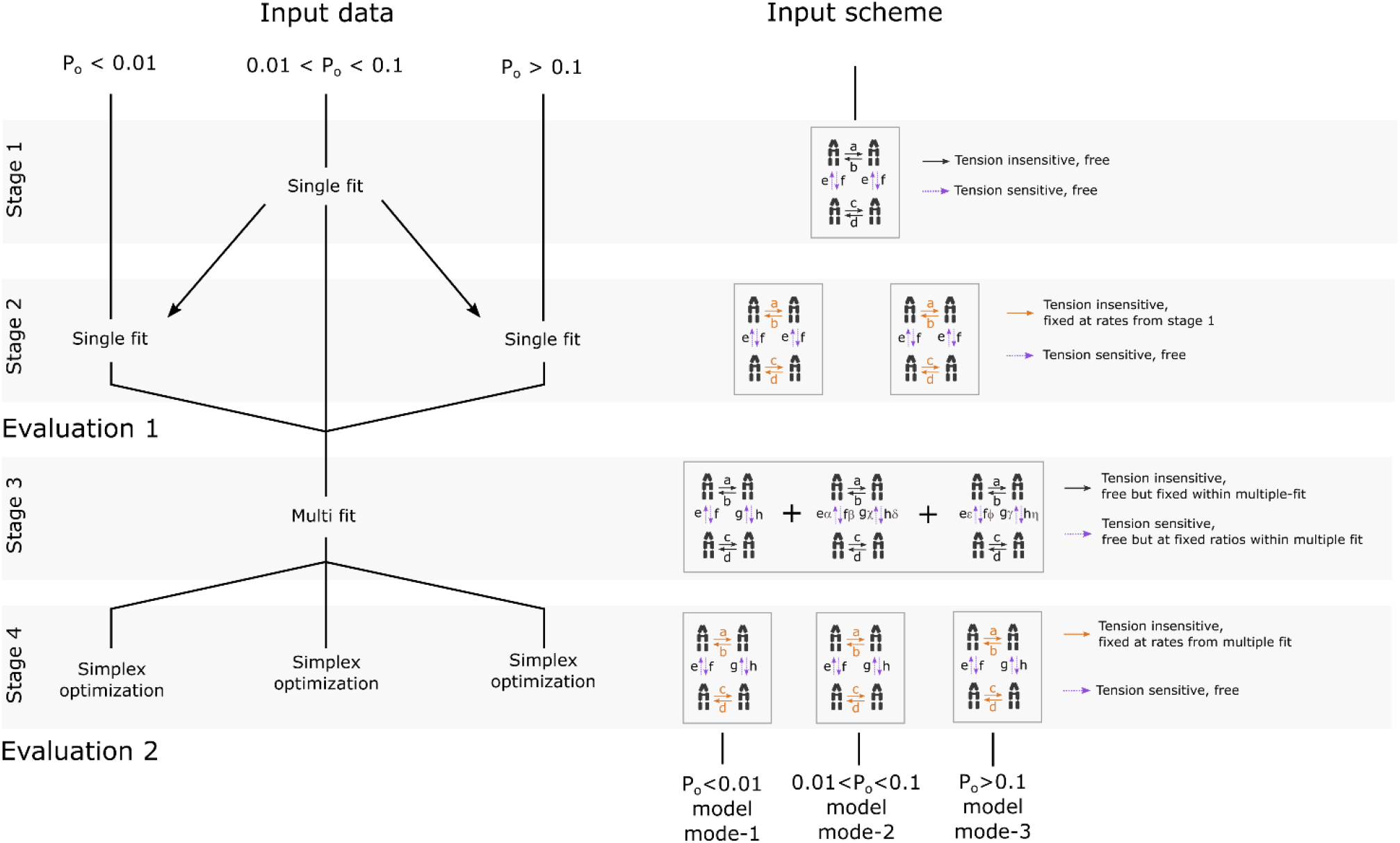
Overview of the fitting procedures used through this study. The fitting protocol takes three dwell-time segments with low, mid, and high P_o_ and a scheme that postulates how specific states are connected. Here, an exemplary four-state scheme with four tension sensitive and four tension-insensitive rates illustrate various aspects of the fitting protocol. Through four stages, the protocol combine information from the three input dwell-time segments to produce three model-modes where only tension sensitive rates differ. Stage 1 is a global fit to the mid-P_O_ segment. In stage 2, tension insensitive-rates derived in stage 1 fit are fixed while tension-sensitive rates are determined from independent fits to low and high P_o_ data. Between stages 2 and 3, results from 50 stage 1-2 fits are evaluated (Fig. 4). Stage 3 simultaneously fit the three input dwell-time data sets to the input scheme using stage 2 information on the ratios that tension sensitive rates vary with between fits from the three datasets (symbolized with Greek letters). Finally, stage 4 uses a simplex optimization strategy where tension-insensitive rates derived from stage 3 are fixed. Tension-sensitive rates from stage 3 are used as initial seeds and optimized to the respective dwell-time data sets. After 50 repetitions of stages 3 and 4, the final models are evaluated as in figure 5.

**Figure S5.**
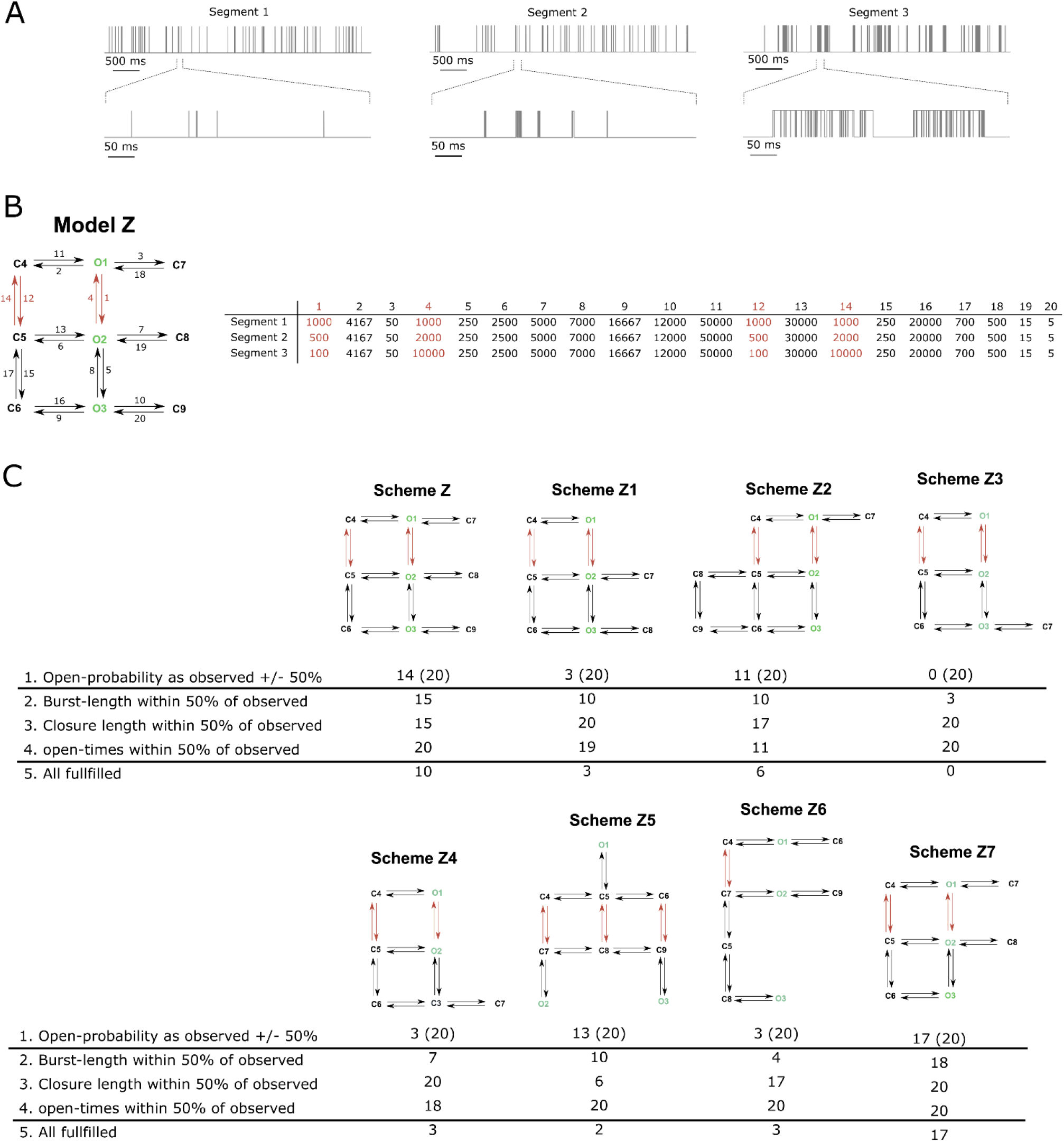
A) Exhibits from three simulated dwell-time segments with increasing channel activity. B) The three dwell-time segments displayed in A were simulated with the basis on Model_Z that have parameters as given in the table. Only four rates (1, 4, 12, and 14) vary between the three segments. Following data simulation, a 20 μs deadtime was imposed. C) Eight schemes assayed through stages 1 and 2 of the *fit-protocol* (Fig. S4) with the three Model_Z dwell-time segments. Each scheme contains open (O) and closed (C) states, and parameters that remain fixed (black) and free (brown) between the three segments. For each scheme, the fit-protocol was repeated twenty times to produce twenty sets of models to account for the three segments. Each repetition was evaluated according to the listed criteria and the number of repetitions fulfilling each criterion given below schemes. The values in parenthesis under criteria 1 indicate the number of fits with P_O_’s that match the initial mid-P_O_ segment; the numbers outside parenthesis represent fits that match P_O_’s for all segments.

**Figure S6.**
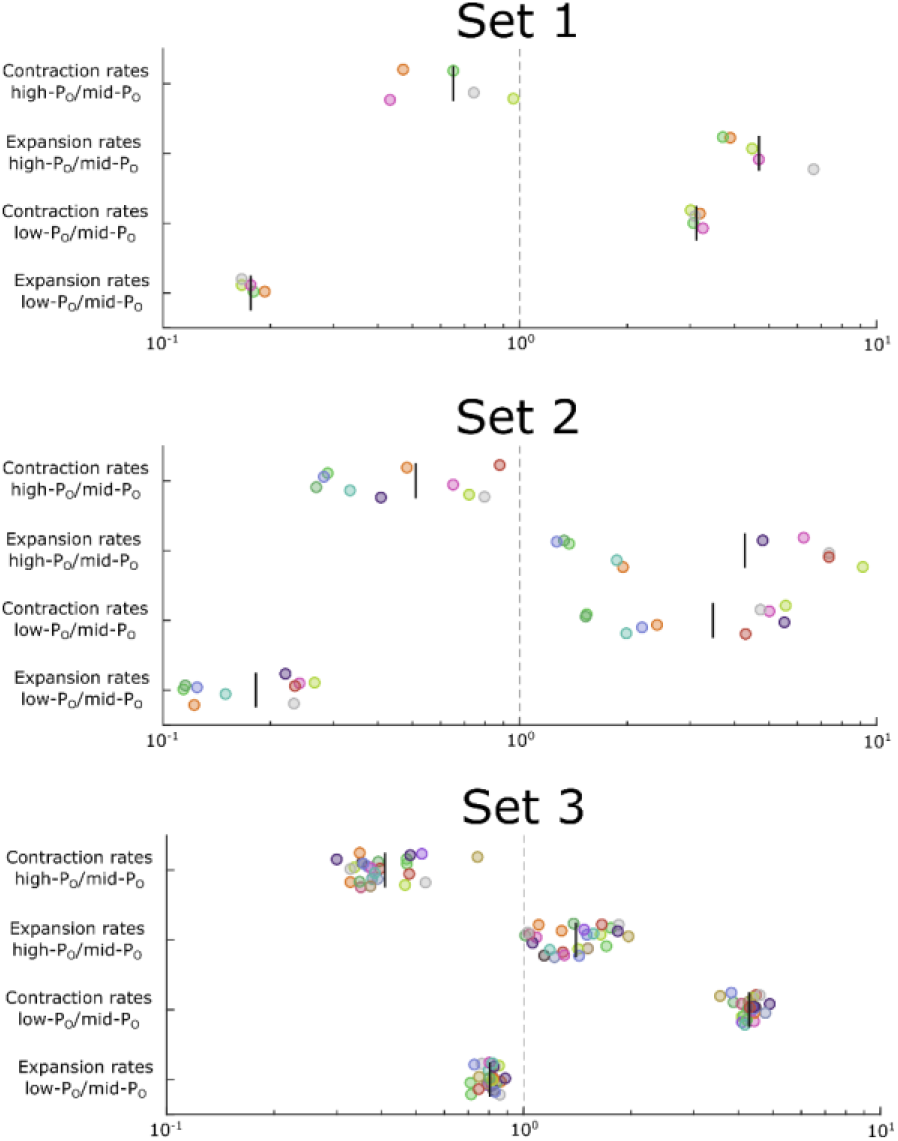
Evaluation of stage 2 Scheme_K demonstrate 5, 11 and 17 models, for dwell-data sets 1, 2 and 3 respectively, that fulfill all criteria. Here the ratio between contraction-rates and expansion-rates for the three dwell-time segments of each set is plotted. Data-points are colored after model, and vertical lines give the mean fold-change values.

**Figure S7.**
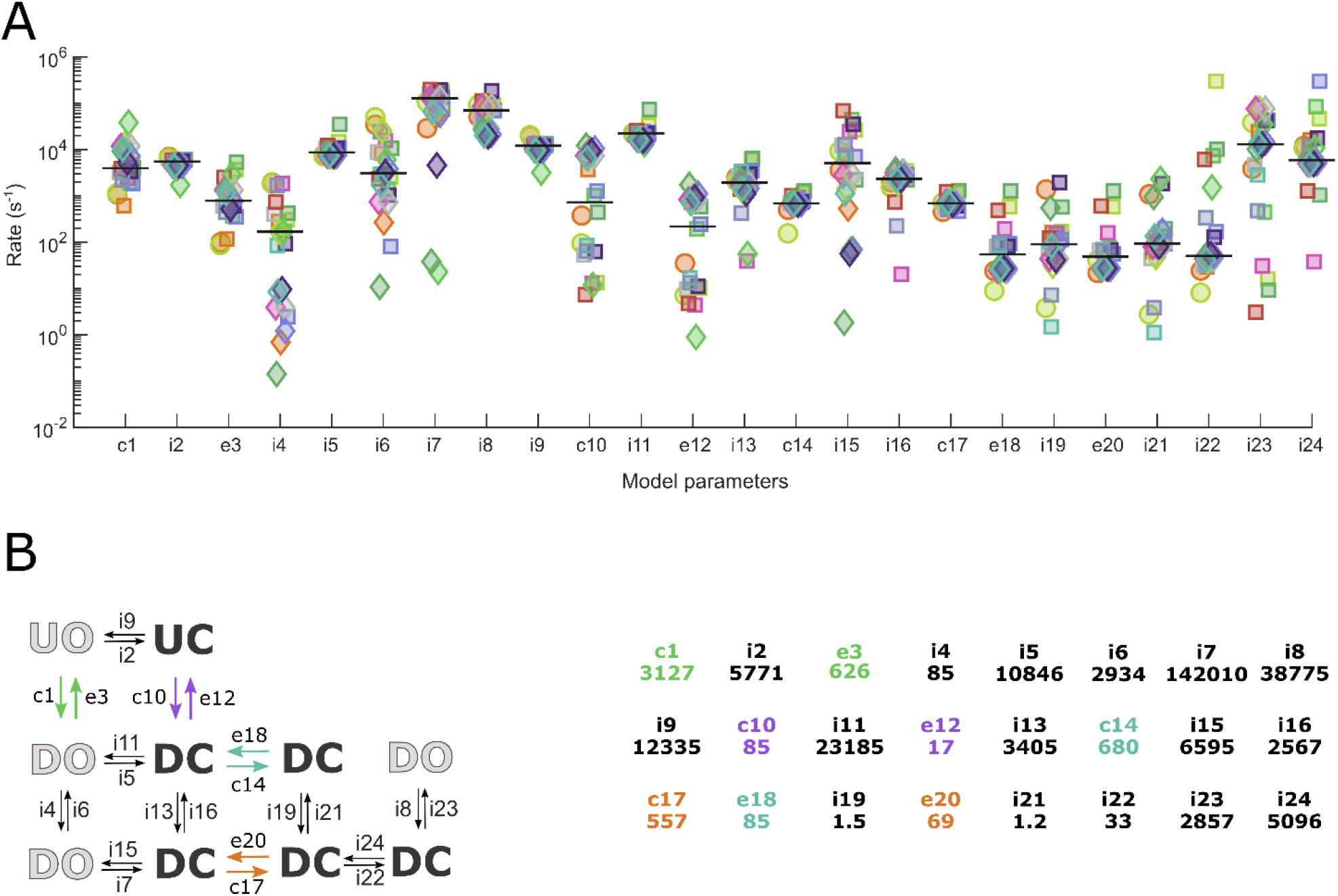
A) Rate constants from the 22 fits of Scheme Kb that fulfill all criteria (Fig 5). These fits are from mid-P_O_ segments, with circles representing set 1, squares set 2 and diamonds set 3, and with symbol colors representing individual fits throughout. The black horizontal lines represent the median value for each parameter. B) The final TREK-2 kinetic model with parameters in the scheme as those given in the table (right). Parameter values in the table are in s^−1^ and labeled as a letter (c = contraction, e = expansion and I = insensitive) followed by a unique number. The final model parameters correspond to the values of the single fit in panel A that have parameters closest to the median values overall.

**Table S1.** Three sets of dwell-times that each include low, mid, and high P_O_ data. Individual segments are color-coded in accordance with Figures 1 and 3.

| Set | $P_o$ | Dwell-times |
| --- | --- | --- |
| 1 | 0.0023 | 11783 |
| 1 | 0.0713 | 11771 |
| 1 | 0.3667 | 15000 |
| 2 | 0.0033 | 11593 |
| 2 | 0.0931 | 24407 |
| 2 | 0.5934 | 25000 |
| 3 | 0.0057 | 10439 |
| 3 | 0.0573 | 13115 |
| 3 | 0.2042 | 12469 |

## References

1. Schmidt, C., et al., Stretch-activated two-pore-domain (K2P) potassium channels in the heart: Focus on atrial fibrillation and heart failure. Prog Biophys Mol Biol, 2017. 130(Pt B): p. 233–243.

2. Talley, E.M., et al., Cns distribution of members of the two-pore-domain (KCNK) potassium channel family. J Neurosci, 2001. 21(19): p. 7491–505.

3. Brohawn, S.G., et al., The mechanosensitive ion channel TRAAK is localized to the mammalian node of Ranvier. Elife, 2019. 8.

4. Kanda, H., et al., TREK-1 and TRAAK Are Principal K+ Channels at the Nodes of Ranvier for Rapid Action Potential Conduction on Mammalian Myelinated Afferent Nerves. Neuron, 2019. 104(5): p. 960-+.

5. Decher, N., A.K. Kiper, and S. Rinne, Stretch-activated potassium currents in the heart: Focus on TREK-1 and arrhythmias. Prog Biophys Mol Biol, 2017. 130(Pt B): p. 223–232.

6. Mathie, A. and E.L. Veale, Two-pore domain potassium channels: potential therapeutic targets for the treatment of pain. Pflugers Arch, 2015. 467(5): p. 931–43.

7. Pereira, V., et al., Role of the TREK2 potassium channel in cold and warm thermosensation and in pain perception. Pain, 2014. 155(12): p. 2534–44.

8. Nedumaran, B., et al., Association of genetic polymorphisms in the pore domains of mechano-gated TREK-1 channel with overactive lower urinary tract symptoms in humans. Neurourol Urodyn, 2019. 38(1): p. 144–150.

9. El Hady, A. and B.B. Machta, Mechanical surface waves accompany action potential propagation. Nat Commun, 2015. 6: p. 6697.

10. Hill, B.C., et al., Laser interferometer measurement of changes in crayfish axon diameter concurrent with action potential. Science, 1977. 196(4288): p. 426–8.

11. Berrier, C., et al., The purified mechanosensitive channel TREK-1 is directly sensitive to membrane tension. J Biol Chem, 2013. 288(38): p. 27307–14.

12. Brohawn, S.G., Z. Su, and R. MacKinnon, Mechanosensitivity is mediated directly by the lipid membrane in TRAAK and TREK1 K+ channels. Proc Natl Acad Sci U S A, 2014. 111(9): p. 3614–9.

13. Clausen, M.V., et al., Asymmetric mechanosensitivity in a eukaryotic ion channel. Proc Natl Acad Sci U S A, 2017. 114(40): p. E8343–E8351.

14. Dong, Y.Y., et al., K2P channel gating mechanisms revealed by structures of TREK-2 and a complex with Prozac. Science, 2015. 347(6227): p. 1256–9.

15. Brohawn, S.G., E.B. Campbell, and R. MacKinnon, Physical mechanism for gating and mechanosensitivity of the human TRAAK K+ channel. Nature, 2014. 516(7529): p. 126–30.

16. Lolicato, M., et al., Transmembrane helix straightening and buckling underlies activation of mechanosensitive and thermosensitive K(2P) channels. Neuron, 2014. 84(6): p. 1198–212.

17. Lolicato, M., et al., K2P2.1 (TREK-1)-activator complexes reveal a cryptic selectivity filter binding site. Nature, 2017. 547(7663): p. 364–368.

18. McClenaghan, C., et al., Polymodal activation of the TREK-2 K2P channel produces structurally distinct open states. J Gen Physiol, 2016. 147(6): p. 497–505.

19. Aryal, P., et al., Bilayer-Mediated Structural Transitions Control Mechanosensitivity of the TREK-2 K2P Channel. Structure, 2017. 25(5): p. 708–718 e2.

20. Markin, V.S. and F. Sachs, Thermodynamics of mechanosensitivity. Phys Biol, 2004. 1(1-2): p. 110–24.

21. Bagriantsev, S.N., et al., Multiple modalities converge on a common gate to control K2P channel function. EMBO J, 2011. 30(17): p. 3594–606.

22. Piechotta, P.L., et al., The pore structure and gating mechanism of K2P channels. EMBO J, 2011. 30(17): p. 3607–19.

23. Voldsgaard Clausen, M., Obtaining transition rates from single-channel data without initial parameter seeding. Channels (Austin), 2020. 14(1): p. 87–97.

24. Goehring, A., et al., Screening and large-scale expression of membrane proteins in mammalian cells for structural studies. Nat Protoc, 2014. 9(11): p. 2574–85.

25. Sondermann, M., et al., High-resolution electrophysiology on a chip: Transient dynamics of alamethicin channel formation. Biochim Biophys Acta, 2006. 1758(4): p. 545–51.

26. Colquhoun, D. and A.G. Hawkes, The Principles of the Stochastic Interpretation of Ion-Channel Mechanisms. In: Sakmann B., Neher E. (eds) Single-Channel Recording., 1995.

27. Hawkes, A.G., A. Jalali, and D. Colquhoun, Asymptotic distributions of apparent open times and shut times in a single channel record allowing for the omission of brief events. Philos Trans R Soc Lond B Biol Sci, 1992. 337(1282): p. 383–404.

28. Hawkes, A.G., A. Jalali, and D. Colquhoun, The Distributions of the Apparent Open Times and Shut Times in a Single Channel Record When Brief Events Cannot Be Detected. Philosophical Transactions of the Royal Society of London Series a-Mathematical Physical and Engineering Sciences, 1990. 332(1627): p. 511–538.

29. Colquhoun, D. and B. Sakmann, Fast events in single-channel currents activated by acetylcholine and its analogues at the frog muscle end-plate. J Physiol, 1985. 369: p. 501–57.

30. Colquhoun, D., et al., How to impose microscopic reversibility in complex reaction mechanisms. Biophys J, 2004. 86(6): p. 3510–8.

31. Harrigan, M.P., et al., Markov modeling reveals novel intracellular modulation of the human TREK-2 selectivity filter. Sci Rep, 2017. 7(1): p. 632.

32. Brennecke, J.T. and B.L. de Groot, Mechanism of Mechanosensitive Gating of the TREK-2 Potassium Channel. Biophys J, 2018. 114(6): p. 1336–1343.

33. Blin, S., et al., Mixing and matching TREK/TRAAK subunits generate heterodimeric K2P channels with unique properties. Proc Natl Acad Sci U S A, 2016. 113(15): p. 4200–5.

34. Lengyel, M., G. Czirjak, and P. Enyedi, Formation of Functional Heterodimers by TREK-1 and TREK-2 Two-pore Domain Potassium Channel Subunits. Journal of Biological Chemistry, 2016. 291(26): p. 13649–13661.

35. Levitz, J., et al., Heterodimerization within the TREK channel subfamily produces a diverse family of highly regulated potassium channels. Proceedings of the National Academy of Sciences of the United States of America, 2016. 113(15): p. 4194–4199.

36. Rudmann, D.G., On-target and off-target-based toxicologic effects. Toxicol Pathol, 2013. 41(2): p. 310–4.

